# *wingless* is a positive regulator of eyespot color patterns in *Bicyclus anynana* butterflies

**DOI:** 10.1101/154500

**Authors:** Nesibe Özsu, Qian Yi Chan, Bin Chen, Mainak Das Gupta, Antónia Monteiro

**Affiliations:** Biological Sciences, National University of Singapore, Singapore 117543; Institute of Entomology and Molecular Biology, Chongqing Normal University, Shapingba, 400047 Chongqing, China; Yale-NUS College, Singapore 138614

**Keywords:** Novel trait, morphogen, transgenesis, RNAi

## Abstract

Eyespot patterns of nymphalid butterflies are an example of a novel trait yet, the developmental origin of eyespots is still not well understood. Several genes have been associated with eyespot development but few have been tested for function. One of these genes is the signaling ligand, *wingless*, which is expressed in the eyespot centers during early pupation and may function in eyespot signaling and color ring differentiation. Here we tested the function of *wingless* in wing and eyespot development by down-regulating it in transgenic *Bicyclus anynana* butterflies via RNAi driven by an inducible heat-shock promoter. Heat-shocks applied during larval and early pupal development led to significant decreases in *wingless* mRNA levels and to decreases in eyespot size and wing size in adult butterflies. We conclude that *wingless* is a positive regulator of eyespot and wing development in *B. anynana* butterflies.

## Introduction

The origin of novel traits remains an outstanding question in evolutionary developmental biology (Hall and Kerney, 2012; Monteiro and Podlaha, 2009; Wagner, 2014). In particular, it is largely unknown how novel traits originate via modifications in development (Wagner, 2015). It has been suggested that novel traits arise when pre-existing genes (True and Carroll, 2002) or larger gene regulatory networks (Monteiro and Das Gupta, 2016) get co-opted into novel parts of the body and function in this novel context to produce the new trait. Thus, understanding trait origins can begin with the identification and functional investigation of key molecular players in trait development.

One example of a morphological novelty is the eyespot, a circular pattern with contrasting color rings, on the wings of butterflies. Comparative data suggests that eyespots originated once within the nymphalid family of butterflies, around 90 million years ago (Oliver et al., 2014; Oliver et al., 2012), likely from simpler colored spots (Oliver et al., 2014). Eyespots appear to serve adaptive roles in both predator avoidance and sexual signaling (Kodandaramaiah, 2011; Oliver et al., 2009; Olofsson et al., 2010; Prudic et al., 2011; Robertson and Monteiro, 2005; Stevens, 2005; Stradling, 1976; Westerman et al., 2014; Westerman et al., 2012) and eyespot number and size are key determinants of butterfly fitness (Ho et al., 2016; Kodandaramaiah, 2011; Prudic et al., 2011; Prudic et al., 2015; Robertson and Monteiro, 2005; Stevens et al., 2007; Westerman et al., 2014; Westerman et al., 2012).

Several genes have been associated with butterfly eyespot development via their eyespot-specific expression (reviewed in Monteiro 2015), however, only a few of these genes have been directly tested for function (Dhungel et al., 2016; Monteiro et al., 2013; Monteiro et al., 2015; Tong et al., 2014; Tong et al., 2012; Zhang and Reed, 2016). *wingless* (*wg*) is one of the genes associated with eyespot development in *Bicyclus anynana* butterflies as Wg protein was visualized in developing eyespot centers at the early pupal stage (Monteiro et al., 2006).

Wg is a signaling ligand involved in multiple aspects of animal development. This includes wing growth and differentiation of melanized spots on the wings of *Drosophila* flies (Sharma, 1973; Sharma and Chopra, 1976; Werner et al., 2010), as well as pigmentation in the silkworm, *Bombyx mori* (Yamaguchi et al., 2013; Zhang et al., 2015). *wg* down-regulation via local electroporation of short interfering RNA (siRNA) showed that *wg* is required for the development of crescent-like melanized markings on the larval epidermis of *B. mori* (Yamaguchi et al., 2013), whereas knock-outs in the same species with CRISPR-Cas9 showed lighter embryo pigmentation effects despite almost complete embryonic lethality (Zhang et al., 2015). On the other hand, spots of dark pigment can be induced by the ectopic expression of *wg* in particular regions of the wings of *Drosophila guttifera* (Werner et al., 2010), and in the larval epidermis of *B. mori* (Yamaguchi et al., 2013), showing the sufficiency of *wg* in generating these patterns. Furthermore, genetic variation in the vicinity of the *wingless* locus controls variation in number of larval markings in *B. mori* silkworms (Yamaguchi et al., 2013). Additionally, a recent study (Koshikawa et al., 2015) showed that a novel enhancer of *wg* is associated with a novel wing color pattern in *Drosophila guttifera* flies. Since evolution in the regulation of *wg* appears to be involved in the origin of novel wing color patterns in flies and lepidoptera, we set out to test *wg* function in eyespot development in butterflies.

Differentiation of the rings in a butterfly eyespot has been hypothesized to result from the action of a morphogen produced in the eyespot center that diffuses to neighboring cells during the early pupal stage (Monteiro et al., 2001; Nijhout, 1980). The morphogen hypothesis is supported by experiments where transplantation of cells from the future eyespot centers induce a complete eyespot in the tissue around the transplant (French and Brakefield, 1995; Monteiro et al., 1997; Nijhout, 1980), and where damage inflicted to these central cells leads to reductions in eyespot size (Brakefield and French, 1995; French and Brakefield, 1992; Monteiro et al., 1997). Although other mechanisms, such as serial induction of the rings, have been proposed for eyespot differentiation (Otaki, 2011), the morphogen hypothesis can most easily explain why central damage can sometimes induce outer rings of color bypassing the induction of the inner rings (Monteiro, 2015).

Both Wg and TGF-β ligands were proposed as candidate morphogens involved in butterfly eyespot formation due to the presence of Wg protein and pSmad protein, a signal transducer of the TGF-β signaling pathway, at the center of the pattern in *B. anynana*, when signaling is known to be taking place (Monteiro et al., 2006). Here we test the function of one of these candidates, *wg*, in eyespot and wing development by down-regulating this gene in independent transgenic lines using a heat-shock inducible wg-RNAi construct, and measuring the effect of this down-regulation on adult eyespot size, wing size, and body size.

## Materials and Methods

### Animal husbandry

Butterflies were reared in climate controlled chambers at 27°C on a 12L: 12D photoperiod, and 80% humidity. Larvae were fed with young corn plants and adults with mashed banana.

### In-situ hybridization

A *wg* riboprobe was synthesized from a *wingless* 558 bp fragment, amplified from cDNA (with primers *wg_*F: 5′ - CCA TGT GGA CCG CTC GCC GC - 3′ and *wg_*R: 5′ - GTG TCG TTG CAG GCA CGC TCG - 3′) and cloned into a pGEMT-Easy vector. For *in situ* hybridization, we used a modified version of the protocol in (Martin and Reed 2014). The sequence of the probe used is provided in Suppl. File 1.

### Making the *wg*-RNAi transgenic lines

A wg-RNAi vector was constructed using the *piggyBac* vector, Pogostick (Chen et al., 2011). Two reverse complementary and complete cDNA sequences of *B. anynana wg* were cloned in opposite direction into the vector. These fold upon each other upon transcription, and initiate the process of RNAi inside the cells. The activation of the RNAi process is controlled temporally by a heat-shock, via the heat-shock promoter from *Heat-shock protein 70 (Hsp70*) from *Drosophila*, which is functional in *Bicyclus* butterflies (Chen et al., 2011; Ramos et al., 2006). Eggs were injected with a mix of the wg-RNAi vector (800 ng/ul in the final concentration), a piggyBac helper plasmid (800 ng/ul), and a small amount of food dye within one hour after being laid, following the protocol of (Ramos et al. 2006). Hatched larvae were placed on a young corn plant and reared to adulthood. Groups of up to five individuals of the same sex were placed in the same cage with the same number of wild-type butterflies of the opposite sex for mating to take place. Their offspring were screened for the expression of green fluorescence in the eyes. Contained within Pogostick is a marker for transformation that contains the gene for *Enhanced green fluorescent protein (Egfp*) driven by a synthetic promoter (*3xP3*) that drives gene expression in the eyes up to adult emergence (Chen et al., 2011; Gupta et al., 2015). Positive individuals were confirmed via PCR with primers specific to the vector and the *wg* sequence inserted into the vector (Clone_R: 5′ - AAC GGC ATA CTG CTC TCG TT - 3′; *wg_*F: 5′ - GTC ATG ATG CCC AAT AC CG - 3′).

### Whole-body heat-shocks

Three independent heat-shock experiments were carried out in this study. In the first experiment heterozygous transgenic and sibling non-transgenic Wt butterflies were reared at 27°C and given two heat-shock pulses, the first heat-shock started at 2pm (~9 h before pupation), whereas the second heat-shock started 12 h later, at 2am (~3 h after pupation). These two time periods were chosen based on previous work that showed a ~8 h delay in the RNAi response following a heat-shock and a loss of the down-regulation effect ~38 h after a single heat-shock performed at 39°C (Chen et al., 2011). The intended goal was to down-regulate *wg* in eyespots from the moment of pupation to around 24 h after pupation, when eyespot ring differentiation is thought to be complete (French and Brakefield, 1995), and Wg protein expression is no longer visible in the eyespot field (Monteiro et al., 2006). Pupae normally pupated between 11 pm and 12 am. Heat-shocks were performed at 39°C for 1.5h (Tong et al., 2014). Similar numbers of transgenic and sibling wild-type butterflies, not exposed to heat-shock, were used as controls. Pre-pupae pupated within the incubator, and the resulting pupae were removed before 2pm the following day. These pupae were later screened for their genotype: Heterozygous wg-RNAi animals with green fluorescence eyes were separated from their wild-type siblings before adult eclosion. The second heat-shock experiment was applied to homozygous transgenic and non-sibling wild-type butterflies of a subsequent generation and followed the same heat-shock conditions as the first experiment (Table 1). The third heat-shock experiment was applied to homozygote individuals of a subsequent generation and consisted of multiple heat-shocks. Homozygous transgenic and wild-type butterflies reared at 27°C were heat-shocked four times a day, at 39°C for 1.5h, with a 6 hour interval, from the beginning of the fifth larval stage till adult eclosion. All heat-shocks were conduced in a Sanyo laboratory incubator oven (MIR152).

### Morphological measurements

Adults were sacrificed by freezing shortly after emergence. Left forewings from female butterflies were carefully cut from the body and imaged using a digital microscope with an attached camera (Leica DMS1000). Pictures were taken using a Leica 0.32X lens at 2.52 magnification. Wings were measured without knowledge of line or treatment identity in Adobe Photoshop. The dorsal forewing Cu1 eyespot of females was selected for measurements as it exhibits minimal developmental plasticity in response to temperature, and is therefore expected to be less responsive to the effects of heat-shocks, as opposed to male dorsal eyespots and ventral eyespots (Monteiro et al., 2015; Prudic et al., 2011). This minimizes confounding effects of heat on eyespot size. Nevertheless, we control for these confounding effects by comparing whether heat-shocked individuals from Wt and transgenic lines respond to the heat-shock in the same way (see statistics below). The following five traits were measured on all dorsal female forewings: the area of the white center, black ring, and gold rings of the Cu1 eyespots, the whole eyespot area obtained by adding the three measurements above, and the whole wing area. Eyespot measurements were done using the ellipse tool to draw the limits of each color ring manually, and using the magic wand tool to select the whole wing area in Adobe Photoshop. Fresh body mass (weight) was measured after the wings were removed from the bodies.

### Real-time PCR

To confirm *wg* knock-down, *wg* mRNA levels were measured before and after the heat-shock treatments by quantitative PCR (qPCR). Wing tissue was dissected from *wg*-transgenic and sibling Wt pre-pupae and early pupae at different time points before and up to 18 h after the first heat-shock, with a 6 hr interval between each time point, and stored in RNAlater solution (Qiagen) at −80°C. The following time points were sampled: At 2 pm before the start of the first heat-shock (BH), 6 h later, 12 h later (and before the 2^nd^ heat-shock), and 18 h later (after both heat-shocks were applied). Animals were at the pre-pupal stage before the first heat-shock (BH) and 6 h after the first heat-shock, and at the early pupal stage 12 h and 18 h after the first heat-shock. Total RNA was extracted from the set of two forewings from each individual using an RNeasy Plus Mini Kit (Qiagen). RNA was treated with RNase-free DNase I (Thermo Scientific) to prevent genomic DNA contamination. Total RNA concentration and purity were measured using NanoDrop 1000 spectrophotometer (Thermo Scientific). Three biological replicates were used per time point and sample type.

Around 200 ng of RNA per sample was reverse-transcribed to cDNA with Reverse-Transcriptase PCR (RT-PCR) using the RevertAid Reverse Transcription Kit (Thermo Scientific). Real-time qPCR was performed with KAPA SYBR^®^ FAST qPCR Kit (KAPA Biosystems) using the Applied Biosystems ABI Prism 7000 Sequence Detection System. Three technical replicates were run for each biological replicate. Average values of technical replicates were used to calculate expression levels of each sample. For each sample, 5 ng of cDNA was quantified. Amplification and quantification of *wg* cDNA levels used the following *wg* primers: *wg_*F: 5′ - CCG AGA GTT CGT TGA CA - 3′; *wg_*R: 5′ - ACC TCG GTA TTG GGC AT -3′, which amplifies a fragments of 246 bp in length. The housekeeping gene *EF1-α* was used as the reference gene for the relative quantification of *wg* expression because expression levels of *EFl-α* were consistent throughout development and showed similar Ct values for tissue samples collected at different developmental times. *EF1-α* primers used were: *EFl-a_*F: 5′ - GTG GGC GTC AAC AAA ATG GA - 3′; *EFl-a_*R: 5′ - TTA GCG GGA GCA AAA ACA ACG AT - 3′, which amplify a 404 bp fragment. Each reaction mixture contained 10 μl of KAPA Master Mix, 0.5 μl of *wg* or *EF1-α* forward primers, 0.5 μl of *wg* or *EF1-α* reverse primers, 8.1 μl of DEPC-treated water and 0.5 μl of cDNA. For a negative control we used DEPC-treated water, in place of cDNA.

The reaction conditions were 95°C for 3 minutes, followed by 40 amplification cycles of 95°C for 30 seconds, 57°C for 30 seconds and 72°C for 30 seconds. Relative quantification of *wg* transcripts was obtained using the 2^-ΔΔCt^ method (Livak and Schmittgen, 2001), transcript expression levels were normalized to the *EF1-a* gene and one sample was used as a calibrator to compare the expression of *wg* transcripts across developmental time points.

### Statistical analysis

Analyses of covariance (ANCOVA) were performed on adult dorsal forewing measurements, with line (*wg*-transgenic vs Wt) and treatment (heat-shock vs control) as fixed variables, family as a random variable, and with wing size as the covariate to normalize eyespot measurements by wing area because eyespot size is normally positively correlated with wing area (Monteiro et al., 2013). The model included all main effects and two-way interactions such as line*family, line*treatment and family*treatment. Levene’s test was used to test homogeneity of variances between the sample groups compared and analyzed, and data transformations in the form of logarithm or other arithmetic functions were conducted as necessary. In data from the second heat-shock experiment, white center, gold ring and total eyespot size from line A were transformed to log10 values. In data from the third heat-shock experiment, black ring area from line A and white center, black ring, gold ring, and total eyespot area from line B were transformed using 1/x^2^ ratio. Estimated means (of eyespot size features) for each group of butterflies, for the same wing size, are plotted in all graphs.

For the *wg* qPCR data, analyses of variance (ANOVA) were used to test for differences in *wg* relative expression levels at the respective time points in wings extracted from *wg-*transgenic and Wt individuals. Logarithmic data transformations were conducted across all data in order to make variances comparable across groups. SPSS statistics, Version 20, was used for all analyses.

## Results

### ln-situ hybridization shows *wingless* is expressed in eyespot centers

To confirm the presence of *wg* expression in eyespot centers in early pupal wings of *B. anynana* we performed *in situ* hybridizations using a riboprobe against *wg* (Suppl. File 1). We visualized *wg* expression in eyespot centers of forewings and hindwings in wing discs of 16, 17, and 24-26 h old pupae as well as expression along the wing margin (Fig. 1), confirming previous work that detected Wg protein in these regions up to 16 hrs (using an antibody against human Wnt1) (Monteiro et al., 2006), and showing that transcripts are present beyond this period.

**Fig. 1.**
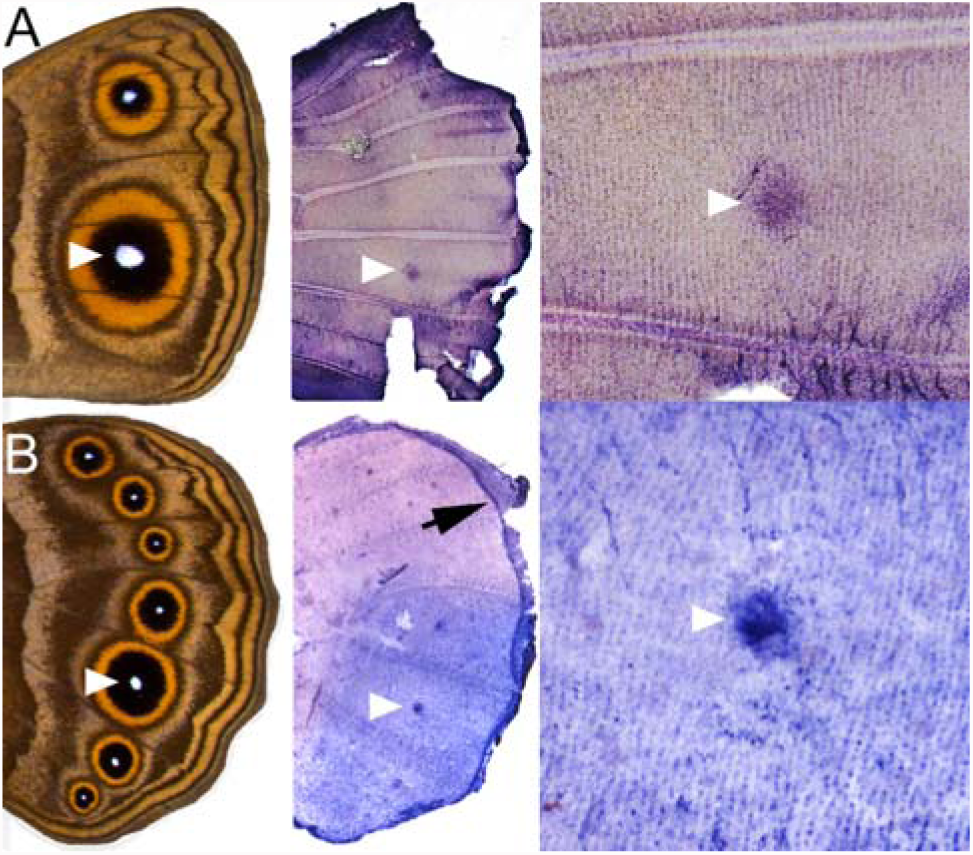
*wg* is expressed in eyespots and in the wing margin. (A) *wg* is expressed in the future eyespot centers (white arrow heads mark the Cu1 eyespots) of a 24-26 h old pupal forewing and (*B*) a 16 h old pupal hindwing, as well as along the wing margin (black arrow).

**Table 1.** Differences between independently conducted heat shock experiments

| Parameters |  |  | Heat shock Experiment I | Heat shock Experiment II | Heat shock Experiment III |
| --- | --- | --- | --- | --- | --- |
| Number of heat shocks per individual |  |  | 2 | 2 | Multiple heat shocks |
| Developmental stage during heat shocks |  |  | Pre-pupae and early pupae | Pre-pupae and early pupae | From 5th larval instar until eclosion |
| Homogeneity of the transgenic butterflies |  |  | Heterozygous | Homozygous | Homozygous |
| Sample size | Line A | Heat shocked | Line A: 57 | Line A: 30 | Line A: 17 |
|  |  |  | Wild-type control: 70 | Wild-type control: 30 | Wild-type control: 31 |
|  |  | Non-heat shocked | Line A: 57 | Line A: 30 | Line A: 27 |
|  |  |  | Wild-type control: 52 | Wild-type control: 30 | Wild-type control: 33 |
|  | Line B | Heat shocked | Line B: 101 | Line B: 30 | Line B: 38 |
|  |  |  | Wild-type control: 54 | Wild-type control: 30 | Wild-type control: 31 |
|  |  | Non-heat shocked | Line B: 58 | Line B: 30 | Line B: 46 |
|  |  |  | Wild-type control: 36 | Wild-type control: 30 | Wild-type control: 33 |
| Data used for |  |  | Morphological measurement and gene expression | Morphological measurement | Morphological measurement |

### Making the transgenic lines

Wild-type embryos were injected with a *wg*-RNAi piggyback based vector (Pogostick) (Chen et al., 2011), as well as a helper plasmid. The *wg*-RNAi construct contains a heat-shock promoter that can be used to induce *wg* knock-down upon delivery of a heat-shock. From a total of 7839 injected embryos, 426 larvae hatched (5% hatching rate), and around 60% of the hatched larvae survived to adult stage. Groups of five emerged adults were crossed with Wt virgins of the opposite sex in separate cages. Offspring from two separate cages (line A and line B) displayed high levels of green fluorescence in their eyes (a marker for transgenesis inserted alongside the *wg* inverted sequences; Fig. S1), indicating independent genomic insertions of the *wg*-RNAi construct. The presence of these insertions in EGFP-expressing individuals was confirmed via PCR on genomic DNA extractions. Adults stopped expressing EGFP in their eyes immediately upon emergence, as previously described for this eye-specific promoter (*3×P3*) in *B. anynana* (Gupta et al., 2015). Five offspring of line A and four offspring of line B were crossed with Wt virgins of the opposite sex in separate mating cages to rear separate families. Approximately half of the offspring in each family had bright green eyes, indicating that line A and line B individuals were likely heterozygous for a single genomic insertion. These mixed *wg*-RNAi transgenic and Wt sibling offspring were used for the first heat-shock experiment (Table 1). A few of these heterozygous EGFP-expressing individuals were subsequently mated with each other and offspring with the brightest eyes (~10%) were selected to set-up homozygous transgenic lines (Chen et al., 2011). Individuals from these subsequent generations all had green fluorescent eyes and were used for the second and third heat-shock experiments (Table 1).

### *wingless* is down regulated in wg RNAi transgenic lines

To examine how a pre-pupal and a pupal heat-shock impacted natural *wg* expression we quantified *wg* expression levels in non-heat-shocked (control) and heat-shocked Wt individuals at four developmental time points, from prior to pupation till approximately 6 h after pupation using qPCR applied to whole forewings. *wg* expression was relatively low in control Wt individuals at the early pre-pupal and early pupal stage, compared to the late pre-pupal stage and 6 h post-pupation (PP) (Fig. 2*A*). Heat-shocked Wt butterflies showed higher *wg* expression relative to controls at 6 h and 18 h after the first heat-shock (Fig. 2*A*), this increase was not statistically significant (F_1,6_ = 2.332, p-value = 0.201 at 6 h and F_1,6_ = 0.288, p-value = 0.620 at 18 h). *wg* gene expression was relatively low at 12 h (right after pupation) in both treatment groups, indicating a natural low expression at this time point.

**Fig. 2.**
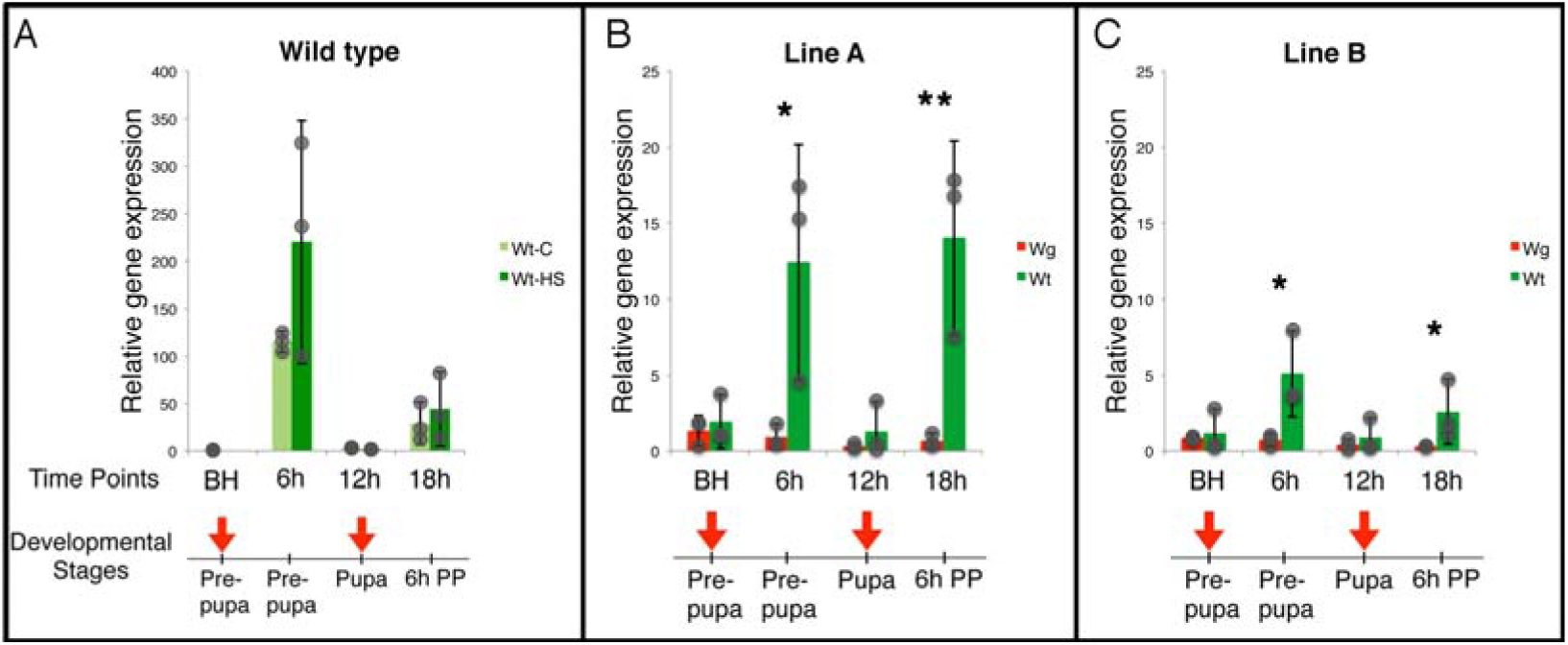
*wg* transcript levels are reduced in wg-RNAi individuals of both lines following one and two heat-shocks. (*A*) *wg* expression (quantified via qPCR) in control (light green bars) and heat-shocked (dark green bars) Wt forewings from the pre-pupal stage to the 6h post-pupal (PP) stage. Heat-shocked Wt butterflies showed comparable levels of *wg* expression relative to Wt controls at 6h and 18h after the first and second heat-shocks, respectively, whereas expression levels were naturally low at the other two time periods. (*B*) In line *A, wg* expression was significantly reduced in *wg*-RNAi wings (red bars) at 6h after the first heat-shock treatment, and at 18h, after the first two treatments, relative to wings of heat-shocked wild-type siblings (green bars). (*C*) In line B, *wg* expression was also significantly reduced in *wg*-RNAi wings at 6h and 18 h after the first heat-shock treatment, relative to wings of heat-shocked wild-type individuals. Arrows indicate the time points of the heat-shock treatments at the pre-pupal and the early pupal stages. Quantification of *wg* mRNA levels at those periods was performed before the heat-shock was applied. Gray dots show the actual data points. Error bars represent 95% confidence intervals of means. * Represents a p-value ≤ 0.05 and ** represents a p-value ≤ 0.01.

To confirm that the heat-shocks were down-regulating *wg* in *wg*-RNAi transgenics, we examined *wg* gene expression in heat-shocked *wg*-RNAi heterozygous individuals and their Wt siblings from both line A and line B. In line A, *wg* expression was significantly down-regulated at 6 h (F_1,6_ = 18.875, p-value = 0.012) and 18 h (F_1,6_ = 46.833, p-value = 0.002) after the first heat-shock relative to wild-type siblings (Fig. 2*B*). Similarly, in line B, *wg* expression in *wg*-RNAi butterflies was also significantly reduced relative to their wild-type siblings at 6 h (F_1,6_ = 18.438, p-value = 0.013) and at 18 h (F_1,6_ = 12.873, p-value = 0.037), after heat-shock treatment (Fig. 2*C*). In addition, there was a large difference in the overall levels of *wg* expression in Wt individuals segregating out of lines A and B at 18 h, after both heat-shock treatments, with wild-type line B individuals displaying lower *wg* levels relative to line A (F_1,6_ = 13.122, p-value = 0.022).

### *wingless* down regulation reduces the size of eyespots

The application of two heat-shocks around pupation led to no changes in wing area (F_1,236_ = 1.079, p-value = 0.300) but led to different responses in eyespot size in *wg*-RNAi transgenic and Wt sibling individuals of line A. Cul dorsal eyespots became reduced in transgenics, relative to non-heat-shocked transgenic controls, while they suffered no change or showed slight increases in size in heat-shocked wild-type sibling butterflies (Fig. 3). This led to a significant interaction between genotype (transgenic and wild-type individuals) and treatment (heat-shock and control) for multiple eyespot area measurements. This interaction was significant for the size of each colored area of scales in an eyespot including the white center (F_1,236_ = 5.163, p-value = 0.024), black disc (F_1,236_ = 4.206, p-value = 0.041) and gold ring (F_1,236_ = 4.279, p-value = 0.040), as well as total eyespot area (F_1,236_ = 4.946, p-value = 0.027) (Fig. 3). However, butterflies from the independently derived and genetically distinct line B didn’t show any statistically significant interactions between genotype and treatment in any of the colored scale areas of forewing Cul eyespots: white center (F_1,249_ = 0.289, p-value = 0.591), black disc (F_1,249_ = 1.549, p-value = 0.215), gold ring (F_1,249_ = 0.056, p-value = 0.814), and combined eyespot area (F_1,249_ = 1.080, p-value = 0.300). These butterflies also did not show any changes in wing area (F_1,249_ = 0.079, p-value = 0.778). The smaller difference observed in levels of *wg* expression between heat-shocked Wt and sibling transgenic individuals of line B (Fig. 2*C*) may explain the weaker eyespot responses to *wg* knockdown in transgenic individuals of this line. For this reason, we conducted two new heat-shock experiments: one where we used homozygous transgenic lines, and kept the heat-shock parameters constant, and one where we used homozygous lines and increased the number and frequency of heat-shocks, starting in the early 5^th^ instar larval stage and ending at adult emergence.

**Fig. 3.**
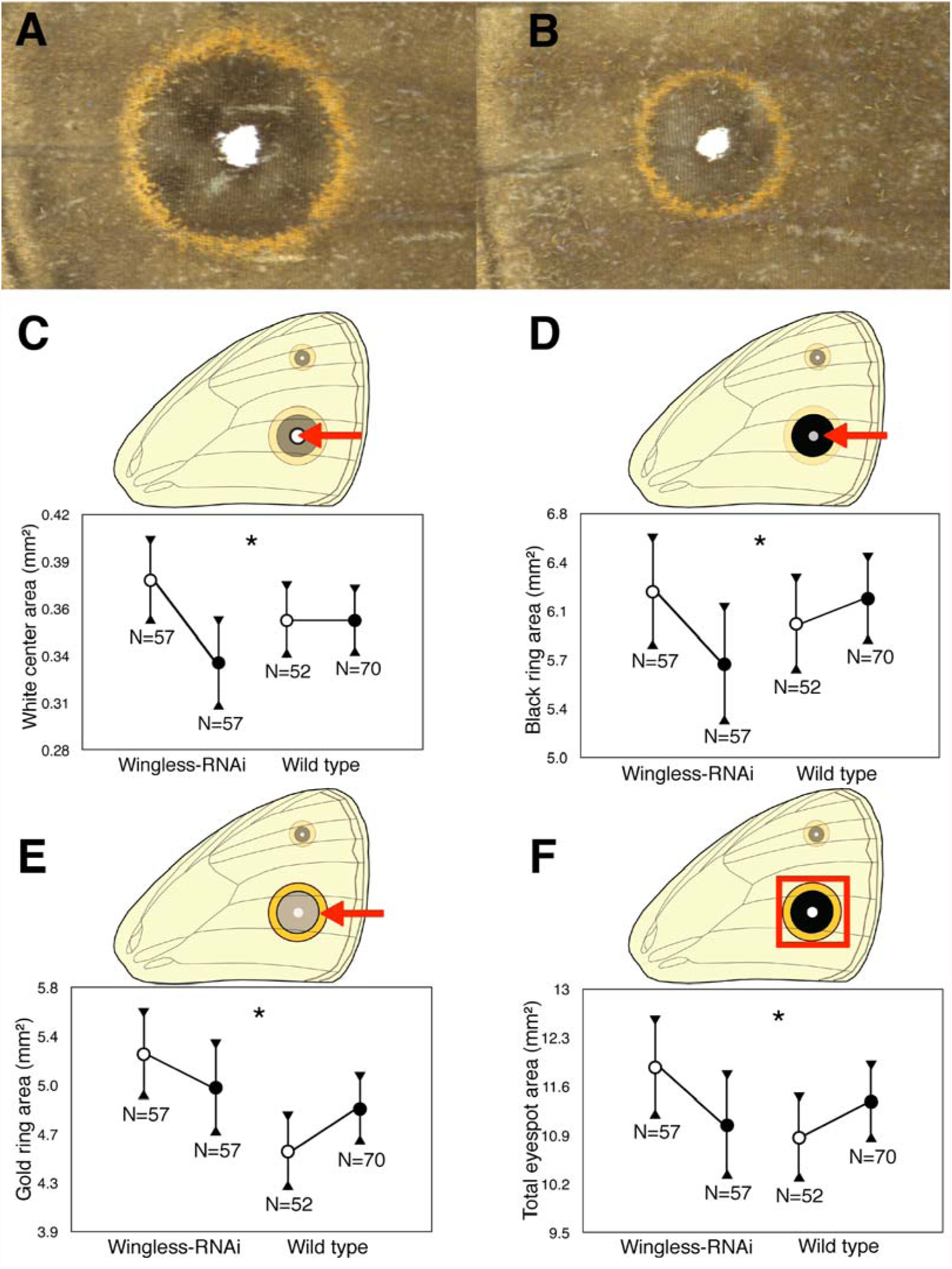
Wt and heterozygous sibling wg-RNAi butterflies show significant differences in their response to two heat-shocks on eyespot size (first heat-shock experiment). (*A*) Representative heat-shocked Wt and (B) heat-shocked *wg*-RNAi transgenic sibling butterflies. Red arrows indicate the Cu1 eyespots measured in this study. Both images are at the same scale. (*C-F*) Area measurements for control (white symbols) and heat-shocked (black symbols) Wt and *wg*-RNAi individuals in the area of the (*C*) white center, (*D*) black ring, (*E*) gold ring and (*F*) total eyespot, with (*) representing a significant interaction between genotype and treatment (p-value ≤ 0.05). Y-axes represent corrected means for each eyespot color ring area, based on values obtained from analyses of covariance on eyespot sizes using wing area as the covariate. Error bars represent 95% confidence intervals of means.

The application of two heat-shocks to homozygous *wg* transgenic and non-sibling Wt butterflies led to similar results as the first experiment using heterozygous individuals. In general, transgenic and non-transgenic individuals responded differently to the heat-shock regarding eyespot size. While heat-shocked transgenic individuals maintained the size of each colored ring, relative to non-heat-shocked individuals, the size of these rings increased in Wt individuals after a heat-shock. This interaction between line and treatment was significant for area of the gold ring (F_1,120_= 5.632, p-value = 0.019) for individuals in line A, and line B (F_1,120_= 7.147, p-value = 0.009). Additionally, p values for the interaction between line and treatment were bordering significance for area of the black ring (F_1,120_ = 3.735, p-value = 0.056), and total eyespot area (F_1,120_= 3.392, p-value = 0.068) in Line A. There were no significant interactions for line and treatment regarding wing area for both lines (Line A: F_1,120_= 0.829, p-value = 0.365; Line B: F_1,120_= 3.296, p-value = 0.072). The use of homozygous individuals of Line B, thus, led to a significant area reduction in one of the color rings, a result not observed with heterozygous individuals. However, the use of non-related individuals, instead of siblings, appears to have reduced the power of this experiment in detecting significant effects of the heat-shock in line A.

### Multiple heat-shocks lead to no effects on eyespot size but strong effects on wing size

Unlike the treatment with two heat-shocks, multiple heat-shocks led to similar eyespot responses in wild-type and *wg*-RNAi individuals of both lines. In general, multiple heat-shocks led to minor changes in the area of all the eyespot color rings relative to wing size in both *wg*-RNAi and Wt individuals (Fig. S2). There were no significant interactions between genotype and treatment in the size of each colored area of scales in the eyespots of line A, including the white center (F_1,108_ = 0.026, p-value = 0.871), black disc (F_1,108_ = 0.092, p-value = 0.763), gold ring (F_1,108_ = 0.000, p-value = 0.987) and overall area (F_1,108_ = 0.023, p-value = 0.880). Similarly, in line B, there was no significant interaction between genotype and treatment in the size of the white center (F_1148_ = 0.308, p-value = 0.580), black disc (F_1,148_ = 0.929, p-value = 0.337), gold ring (F_1,148_ = 2.333, p-value = 0.129), and overall eyespot size (F_1,148_ = 1269, p-value = 0.262). Performing the more extensive series of heat-shocks, however, led to strong effects on wing size (Fig.4), but not on body size (F_1,20_ = 1.864, p-value = 0.189) (Fig. S3). This effect on wing size, where *wg*-RNAi and Wt individuals responded differently to the heat-shocks was not previously observed with the more restrictive pre-pupal/early pupal heat-shocks. Heat-shocking *wg*-RNAi individuals of line A led to a significant reduction in wing size, whereas heat-shocking Wt individuals led to no changes in wing size (line and treatment interaction: F_(1,148)_= 12.657, p-value = 0.001) (Fig. 4). This was also observed in line B (line and treatment interaction: F_(1,148)_ = 11.995, p-value = 0.001) (Fig. 4).

**Fig. 4.**
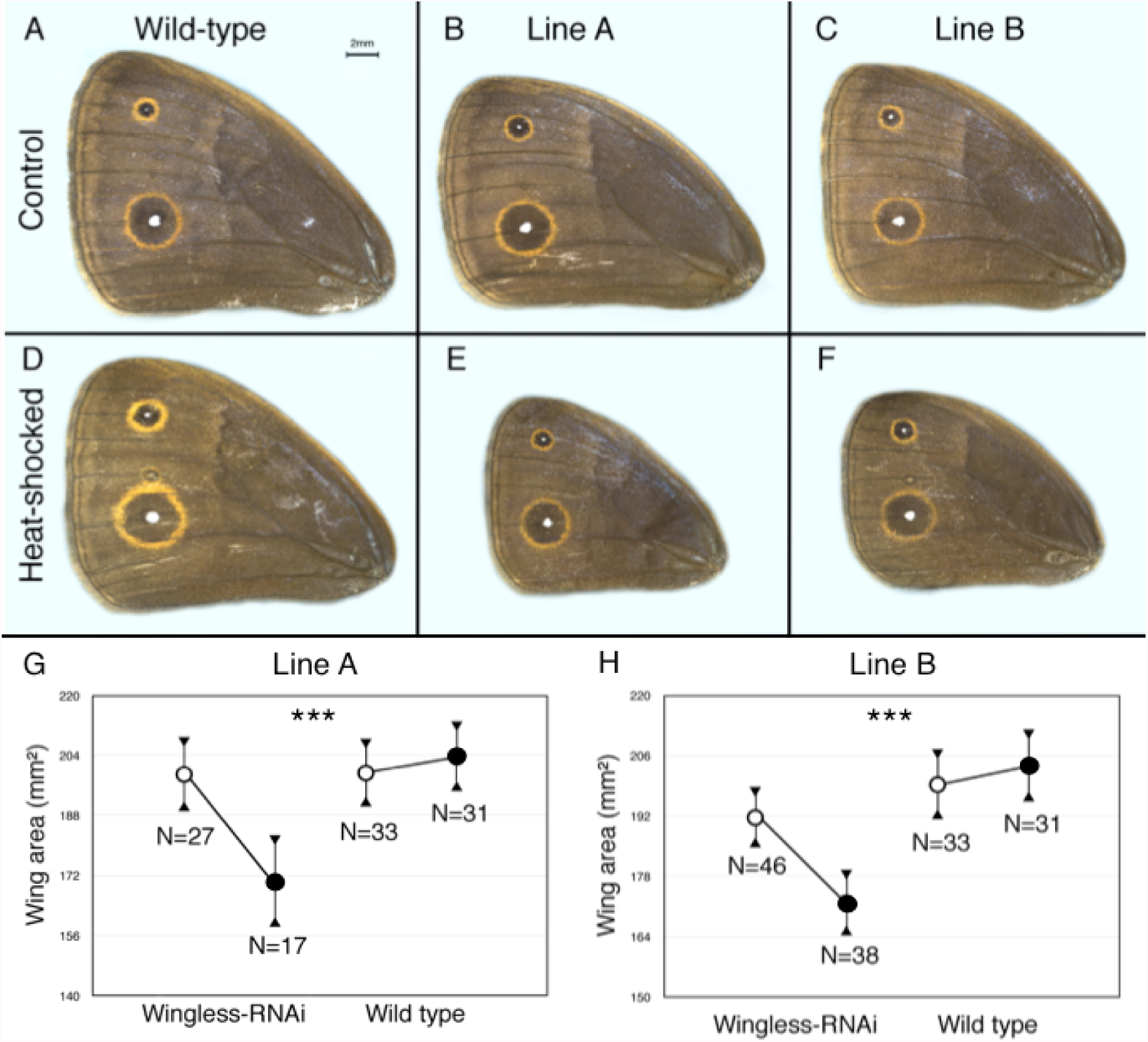
Multiple heat-shocks reduce wing size in homozygous wg-RNAi butterflies of lines A and B but not in wild-type butterflies (third heat-shock experiment). (*A-F*) Representative dorsal forewings of (*A*) control Wt, (*B*) line A and (C) line B individuals, and (D) heat-shocked Wt, (*E*) line A and (*F*) line B individuals. All images are at the same scale (scale bar in A represents 2mm). (*G,H*) Wing area measurements for control (white symbols) and heat-shocked (black symbols) *w*g-RNAi and Wt individuals of (*G*) line A and (*H*) line B. (***) Represents a significant interaction between line and treatment with p-value ≤ 0.001. Y-axes represent the total wing area. Error bars represent 95% confidence intervals of means.

## Discussion

In this study we tested the function of a signaling ligand, *wingless*, in eyespot development using transgenic butterflies carrying a heat-inducible *wg*-RNAi construct. We first showed that *wg* expression was successfully knocked down, albeit to different degrees, in two genetically independent transgenic lines, relative to wild-type sibling butterflies, not containing the transgene. This down-regulation of *wg* led to significant reductions in the size of Cu1 forewing eyespots, for wings of comparable size, indicating that *wg* is a positive regulator of eyespot development in butterflies. Interestingly, our two independently derived transgenic lines had either different endogenous *wg* levels or different sensitivities to the heat-shock, which led to variation in *wg* levels after the heat-shock. More accentuated differences in *wg* levels between heat-shocked and control individuals were found in line A, and less marked differences between treatments in line B. The extent of *wg* variation before and after treatment within a line correlated with the extent of eyespot size variation following heat-shock for each of the lines. In particular, the area of all three color rings was more readily altered in line A (in the first and the second heat-shock experiments), whereas only the area of the outer gold ring was altered in line B (in the second heat-shock experiment).

Reduction of *wg* mRNA levels may be affecting the differentiation of the eyespot rings via changes in a putative Wg protein gradient. If *wg* transcription in the eyespot centers leads to a gradient of Wg protein, diffusing from the central cells to the surrounding cells (Fig. 5), then stronger or weaker modulations in the height and shape of that gradient, could lead to the observed phenotypes (Fig. 5). While the existence of long-range gradients of Wg signaling is currently controversial in *Drosophila* (Alexandre et al., 2014; Martinez Arias, 2003; Strigini and Cohen, 2000), butterfly eyespots may provide an alternative model system to test these ideas in future.

**Fig. 5.**
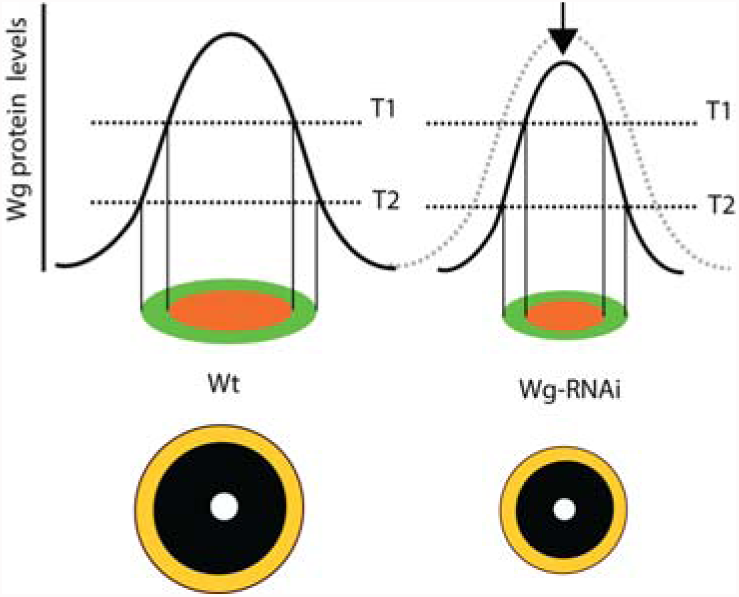
Classic gradient model that can explain how *wg* down-regulation affects the differentiation of the eyespot rings. Differentiation of the rings in a butterfly eyespot could involve a Wg protein gradient (black curved lines) where the protein is produced in the eyespot centers and diffuses to neighboring cells. Threshold responses to that protein gradient could determine the area of the black (T1) and gold (T2) color rings via the activation of intermediate tier genes such as *Distal-less* and *spalt* (red) and *engrailed* (green) (Brunetti et al., 2001). Down-regulation of *wg* (black arrow) alters the area of the color rings in an eyespot, while the thresholds of response to Wg protein remain constant.

The timing of *wingless* expression, measured via *in situ* hybridizations, was found to be extended relative to a previous study that examined *wg* expression at the protein level using cross-reactive antibodies (Monteiro et al., 2006). The previous study showed that Wg proteins were found in the eyespot field (primarily in the center) between 10.5 h and 16 h after pupation, whereas beyond this point, Wg proteins were found at levels below background levels in the eyespot center. Older pupal wings (>24hrs old), however, were not studied (Monteiro et al., 2006). Here, *wg* expression was visualized at the mRNA level in the developing eyespot centers at 16 h and at 22-24 h after pupation. The timing of both mRNA and protein expression fits data from previous experiments where damage applied to the signaling eyespots centers stops having an effect on eyespot size after 24 hrs (French and Brakefield, 1995). However, the reason why Wg protein stops being detected in the eyespot centers after 16 hrs is unknown, and may be due to post-transcriptional regulatory processes not investigated here.

Our results are consistent with the function of *wg* in the development of wing color spots in *D. guttifera* (Koshikawa et al., 2015; Werner et al., 2010) and melanized markings on the larval epidermis in *B. moni* (Yamaguchi et al., 2013), suggesting a conserved role for *wg* in color patterning the integument of flies, moths, and butterflies in the eyespot centers. While these color patterns are not considered homologous, they could be sharing a conserved signaling process for their differentiation.

The current study also demonstrated that *wg* is a positive regulator of wing growth in butterflies similarly to findings in other insects. *wg* down-regulation in butterflies throughout the last (5^th^ larval) instar, as well as throughout pre-pupal and pupal development, led to a significant reduction in wing size in both *wg*-RNAi lines, *wg’s* function in wing growth was initially demonstrated in *D. melanogaster* where frequent occurrences of wingless and haltere-defective fruit flies led to the isolation of the gene (Sharma, 1973; Sharma and Chopra, 1976; Swarup and Verheyen, 2012). *wg* is expressed along the wing margin of larval, pre-pupal, and pupal wing discs in *Drosophila* flies where it promotes wing growth (Couso et al., 1994; Phillips and Whittle, 1993). The same pattern of *wg* expression is observed in *B. anynana* larval (Monteiro et al., 2006) and pupal wings (Fig. 1) as well as larval wings of multiple other butterflies and moths (Carroll et al., 1994; Kango-Singh et al., 2001; Martin and Reed, 2010; Monteiro et al., 2006). Deficiency in *wg* receptors inhibits the development of the wing field (Chen and Struhl, 1999), whereas ectopic expression of *wg* induces overgrowth of wing discs during larval development (Neumann and Cohen, 1997). Levels of *wg* expression are associated with wing length in polymorphic planthoppers, and *wg* RNAi individuals developed significantly shorter and deformed wings (Yu et al., 2014). Lesions in the *wg* gene found in natural populations of Apollo butterflies after a bottleneck were proposed to lead to a high frequency of reduced and deformed wings in individuals of this population (Lukasiewicz et al., 2016). These studies all show that *wg* is required for normal wing growth (Swarup and Verheyen, 2012). Since *wg* expression in *B. anynana* was not completely shut down but merely down-regulated in this study, a lower expression of *wg* in the wing tissues of heat-shocked *wg*-RNAi butterflies led to the development of smaller wings.

Surprisingly, eyespots in *wg*-RNAi and Wt butterflies were affected to the same extent after multiple heat-shocks, i.e., wings of *wg*-RNAi butterflies became significantly smaller but eyespot size scaled down in perfect proportion, rather than disproportionately, with wing size (Fig. S2). It is unclear what factors caused this pattern, but mechanisms of eyespot size plasticity could be playing a role. The eyespots of *B. anynana* are particularly sensitive to ambient temperatures during the wandering stage of late larval development (Monteiro et al., 2015). High temperatures (27°C) during this stage lead to high ecdysteroid titers, which in turn lead to large eyespots (Monteiro et al., 2015). Our multiple heat-shock experiment comprised the wandering stage of development, whereas the late pre-pupal and early pupal heat-shock happened after this stage. It is possible that one of the genes that leads to larger eyespots in response to ambient temperature is *wg*. The positive effect of temperature on *wg* expression could potentially override its negative effect via endogenous *wg* down-regulation leading to relatively proportioned sized eyespots. Interestingly, a connection between the same ecdysteroid and *wg* was observed in *B. mori* larval epidermis where raised ecdysteroid titers at the end of each molt activate *wg* expression in the area of the melanic spots (Yamaguchi et al., 2013).

Recent studies showed that *wg* and *WntA*, another Wnt protein family member, are expressed along anterior-posterior stripes in larval wing discs across multiple species of butterflies (Carroll et al., 1994; Gallant et al., 2014; Martin et al., 2012; Martin and Reed, 2010, 2014). Interestingly, *wg* was found associated with the basal, central and marginal stripe patterns in moths and butterflies (Martin and Reed, 2010), and *WntA* was proposed to play a role in organizing the basal, central, and marginal symmetry systems (Martin and Reed, 2014). Linkage mapping, gene expression, and functional studies using injections of small molecules, heparin and dextran sulfate, that can bind Wnt molecules (as well as other ligands) to enhance their diffusion (Binari et al., 1997; Yan and Lin, 2009), all suggested that *WntA* is associated with the differentiation of anterior-posterior stripes in several butterfly species, including *Euphdryas chalcedona, Junonia coenia, Heliconius* and *Limenitis* butterflies (Gallant et al., 2014; Martin et al., 2012; Martin and Reed, 2014). Here, we show that *B. anynana* eyespots, belonging to the border symmetry system, are in fact using *wg* signaling in the development and differentiation of their color rings. This works constitutes the first functional demonstration that a *Wnt* family member is involved in wing pattern development in butterflies.

Future work should examine whether *wg* ectopic expression would be sufficient to induce an eyespot color pattern in butterflies. This would be necessary to show that the recruitment of this gene to the eyespot centers helped in the origination of a morphological novelty.

## Acknowledgements

We thank Kathy F.Y Su and K.J. Tan for creating *Drosophila* transgenic flies, which tested our original construct, and Ling L. Sheng and Heidi Connahs for performing *in situ* hybridizations with the *wg* probe on embryos. This work was supported by the Singapore Ministry of Education Awards MOE2014-T2-1-146, and MOE R-154-000-602-112 to AM, NUS award R154-000-587-133 to AM, and the Singapore International Graduate Award (SINGA) fellowship from A*STAR to NO.

## Supplemental Figures

**Fig. S1.**
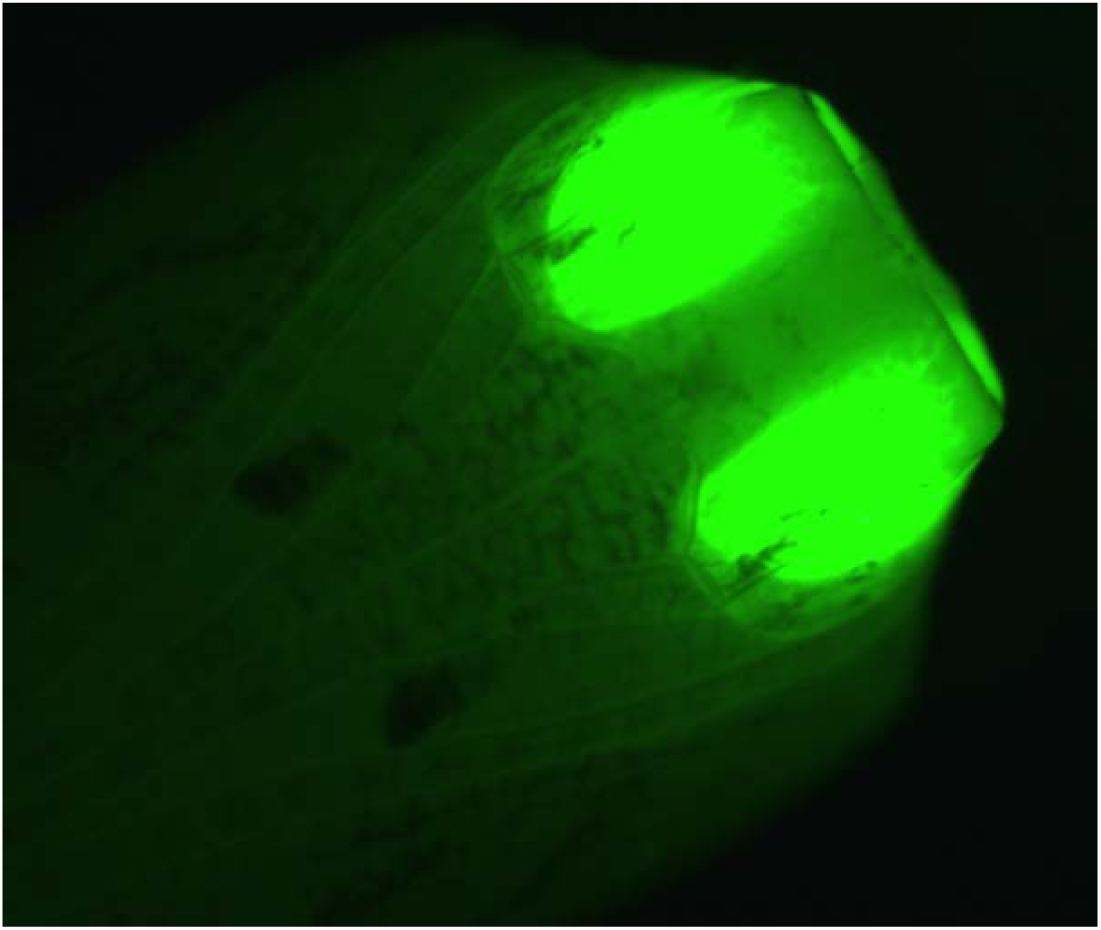
Transgenic *Bicyclus anynana wg*-RNAi line A pupa with green fluorescent eyes.

**Fig. S2.**
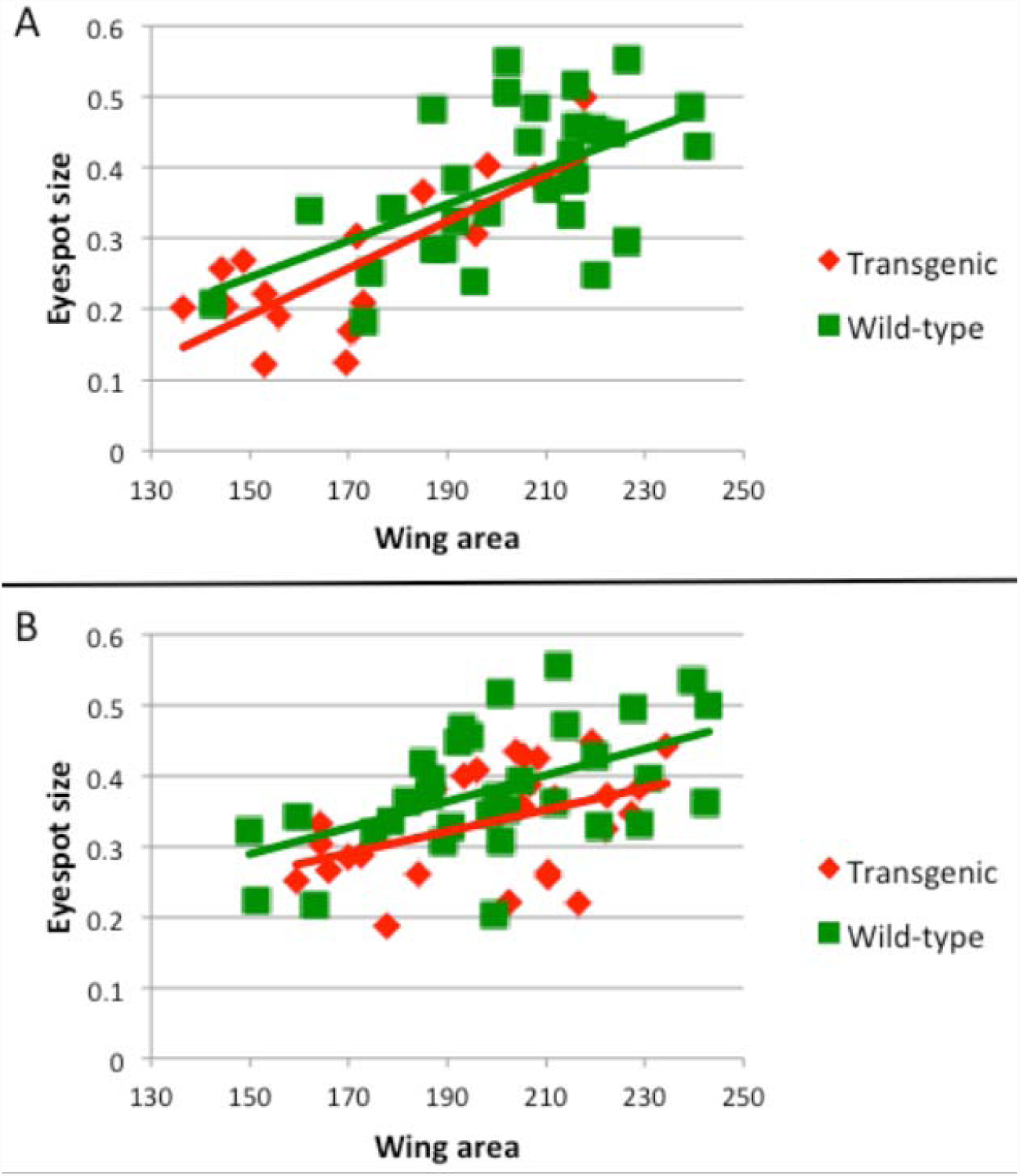
Allometric relationship between eyespot size (white eyespot center) and wing area after the multiple heat-shock treatment. (A) Heat-shocked wild-type and transgenic butterflies of line A. Eyespots are reduced in size in proportion to wing size. (B) Non heat-shocked (control) wild-type and transgenic butterflies of line A.

**Fig. S3.**
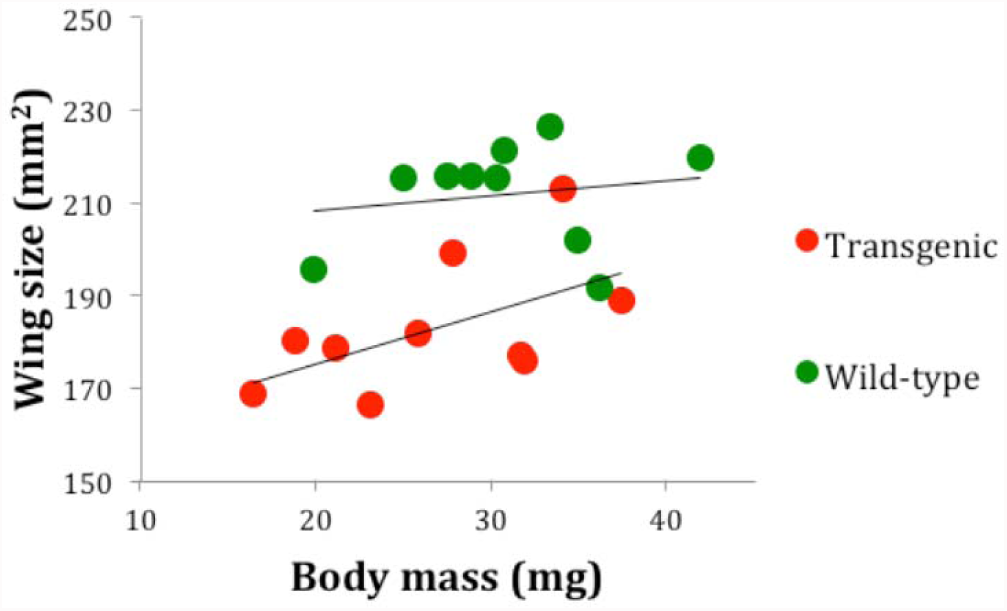
Allometric relationship between wing size and body mass of a random sample of heat-shocked wild-type and transgenic butterflies of line B. Transgenic and wild-type butterflies have different wing sizes but they do not differ in body mass, indicating that *wg* down-regulation has a wing-specific effect.

## Supplementary File 1 – Sequence of *wingless* probe used for the *in situ* hybridizations

CCATNTGGACCGCTCGNCGCACCGCGCGCGNGCCGCCGCCGCCGCCAACGTGAGGGTCTGGAAATGGGGCGGGTGCAGCGACAACATCGGCTTCGGCTTCAAGTTCAGCCGNGANTTCGTTGACACCGGGGAAAGGGGCAAGACGCTTAGGGAGAAGATGAACTTGCACAACAATGAGGCCGGCAGGATGCACGTGCAAACGGAGATGCGCCAGGAGTGCAAGTGCCACGGTATGTCTGGGTCCTGCACGGTGAAGACGTGCTGGATGAGGCTGCCGACGTTCCGGTCTGTAGGCGACGCCCTGAAAGACAGCTTCGACGGGGCGTCGCGGGTCATGATGCCCAATACCGAGGTGGAGGCGCCGTCGCAGAGGAACGACGCCGCACCTCACAGGGTCCCGCGCCGTGACCGCTACAGGTTCCAACTTCGGCCGCACAACCCTGACCACAAAACACCCGGGGTCAAGGACCTTGTATACTTGGAATCTTCACCAGGTTTCTGC

